# Antiviral defence systems in the rumen microbiome

**DOI:** 10.1101/2024.04.04.588093

**Authors:** Johan S. Sáenz, Bibiana Rios-Galicia, Jana Seifert

## Abstract

The continuous interaction between phages and their respective hosts has resulted in the evolution of multiple bacterial immune mechanism. However, the diversity and prevalence of antiviral defence systems in complex communities is still unknown. We therefore investigated the diversity and abundance of viral defence systems in 3038 high-quality bacterial and archaeal genomes from the rumen. In total, 14,241 defence systems and 31,948 antiviral-related genes were identified. Those genes represented 114 unique system types grouped into 49 families. We observed a high prevalence of defence systems in the genomes. However, the number of defence systems, defence system families and systems density varied widely from genome to genome. Additionally, the number of defence system correlated positively with the number of defence systems families and the genome size. Restriction modification, Abi and cas system families were the most common, but many rare systems were present in only 1% of the genomes. Antiviral defence systems are prevalent and diverse in the rumen, but only a few are dominant, indicating that most systems are rarely present. However, the collection of systems throughout the rumen may represent a pool of mechanisms that can be shared by different members of the community and modulate the phage-host interaction.

**Importance:** Phages may act antagonistically at the cell level but have a mutualistic interaction at the microbiome level. This interaction shapes the structure of microbial communities and is mainly driven by defence mechanism. However, the diversity of such mechanism is larger than previously thought. Because of that, we described the abundance and diversity of antiviral defence system of a collection of genomes, from the rumen. While defence mechanisms seem to be prevalent among bacteria and archaea, only a few were common. This suggests that most of these defence mechanisms are not present in many rumen microbes but could be shared among different members of the microbial community. This aligned with the ’pan-immune system’ model, which appears to be common across different environments.

## 1. Introduction

It is estimated that demand for meat and milk will rise between 70-80% of current production by 2050 (1). Part of the current needs are met by raising cattle, which account for almost one third of all domesticated ruminants (2). However, sustainable beef and dairy production is challenged by economic and political issues, environmental factors, feed efficiency, and animal and human health (3, 4). Recent studies have shown that some of these problems are linked to the rumen microbiome.

The massive use of sequencing technologies has shown that the rumen is a complex ecosystem consisting of bacteria, archaea, protozoa, fungi, and viruses (5–7). This group of organisms contribute to the metabolism of complex plant materials and the production of volatile fatty acids (VFAs), which provide most of the energy required by the animal. Rumen bacteria and archaea have been associated with feed efficiency, methane production, animal health and fat and protein content of milk (8–11). Similarly, it has been shown that anaerobic fungi play a crucial role in the efficient degradation of fiber through enzymatic and physical processes (12). Rumen viruses are diverse, organism-specific, predominantly lytic, carry diverse auxiliary metabolic genes and have the potential to modulate nutrients recycling, fiber degradation and methanogenesis (13–16). In addition, dietary changes or perturbations can modulate viral richness and diversity, which has been shown to affect microbial metabolism (16).

Phages shape bacterial and archaeal populations and are a reservoir of accessory genes that facilitate the mobility of genetic material (17). The continuous interaction between phages and their respective hosts has resulted in the evolution of bacterial immune mechanism and phage counter-defence strategies (18, 19). This arms-race is a major driving force of molecular innovation and genetic diversification. Defence mechanism in prokaryotes are diverse and can be divided between innate and adaptive immune systems (20). The prevalence of defence mechanisms ranges from zero to several systems between cells. Defence systems can be compose by single or multiple genes, mainly associated with nucleases, helicases, proteases, kinases, ATPases and reverse transcriptases (21, 22). Antiviral defence mechanism are classify mainly in three groups: nucleic acid degradation, abortive infection and inhibition of DNA/RNA synthesis, but the mechanism of the majority of systems remains unknow (23). Moreover, the effective immunity against phages seems to be driven by within-cell (24), presence of single or multiple system, and pan-genome dynamics (20). Previous studies have shown that rumen bacteria harbour multiple CRISPR-Cas and RM systems (14, 25), which can be associated with the modulation of phage-host interaction. However, a detail description of the diversity and prevalence of antiviral defence systems in the rumen is lacking. We explored the prevalence and diversity of antiviral defence mechanisms in 3038 high quality genomes from the rumen. This study shows that the repertoire of defence mechanisms in the rumen is broad and that the pan-immunity is mainly dominated by innate immune mechanisms.

## 2. Methods

### 2.1. Data collection and dataset description

In total 5990 genomes from the rumen microbiome were collected for the present study. Briefly, 5588 metagenomes assembled genomes (MAGs) from the MGnify (https://www.ebi.ac.uk/about/news/updates-from-data-resources/cow-rumen-catalogue-v10-released-mgnify/) cow rumen v1.0 MAG catalogue and 402 genomes of isolates from the Hungate 1000 catalogue of reference genomes were obtained (5). MAGs dataset is based on data collected from European cattle (26) and African cattle (27).

### 2.2. Quality control of MAGs

CheckM2 (28) was used to estimate the quality of the genomes and only the high-quality genomes with >90% completeness and <5% contamination were kept for further analysis. Additionally, the N50, coding density, gene length, genome size and GC content of the genomes were calculated using CheckM2. The genomes were taxonomically classified using GTDB-tk v2.1 (29) and the GTDB database R207_v2 (30).

### 2.3. Antiviral defence prediction

Antiviral defence mechanisms were identified from all genomes using PADLOC v1.1 (22) and the PADLOC-DB. Systems labelled as other due to incomplete detection of essential genes were discarded. Defence systems were grouped by families using the PADLOC-DB metadata (https://github.com/padlocbio/padloc-db/blob/master/sys_meta.txt) and the references to relevant literature (https://github.com/padlocbio/padloc-db/blob/master/system_info.md). The density of defence systems per genome was calculated as the total defence systems per genome divided by the genome size (kbp). CRISPR-Cas genes and arrays were detected using CRISPRCasTyper v1.6 (31). All predicted CRISPR spacers from complete or incomplete CRISPR-Cas arrays were kept. Viral genomes from the rumen virome database (RVD) were matched against CRISPR spacers using Blast (32), only those with >95% identity, <= 1 mismatch and <0.0004 e-value were considered spacer to protospacer matches. Host-virus interaction network showing the spacer-protospacer match was created using Cytoscape (33).

### 2.4. Prophages detection and identification

Collected genomes were searched for prophages using Virsorter2 v2.2.3 (34). Only contigs longer than >10,000 were used. All contigs identified as prophages were kept for further analysis. Prophages were quality-checked using checkV (35). Prophages with 0 viral genes and >1 host gene were discarded and prophages with 0 viral genes and 0 host genes were kept. A prophage was considered complete if it was predicted as such by checkV.

### 2.5. Phylogenetic placement

Phylogenetic tree of the whole 3038 genomes was created using Phylophlan (36) and 400 universal marker genes, with --medium diversity and --accurate options. Phylogenetic relation among species of *Butyrivibrio*, Ca. *Limimorpha*, *Prevotella* and *Ruminococcus* was calculated using a subset of genes present in at least 90% of all genomes (core genes) for each *Genus*. *Butyrivibrio*, *Prevotella* and *Ruminococcus* were selected due to their key role in the rumen and high prevalence of defence systems, while Ca. *Limimorpha due to its novelty*. Core genes were extracted, aligned, and concatenated using the tool: concatenated-gene-alignment-fasta from Anvi’o v 7.1 (37). The extracted genes (55 of *Butyrivibrio*, 71 of *Limimorpha*, 13 of *Prevotella* and 46 of *Ruminococcus*) were used to observe phylogenetic relatedness. Phylogenetic trees were inferred by maximum-likelihood using the concatenate of core genes of each genome with FastTree2 (38). Taxonomic information, number of defence families and numbers of prophages were displayed in a phylogenetic tree using iTOL (39).

### 2.6. Data wrangling and data availability

All data wrangling and statistical analysis were performed in R v4.1.2 (40) and the packages “tidyverse” (41), “data.table”, “pheatmap”, “ggdist”, “patchwork”, “ggtext”, and “viridis” were used. Medians across groups were compared using the nonparametric Kruskal-Wallis test (*kruskall.test* function). Additionally, correlations were calculated using the Spearman’s rank correlation coefficient (*cor.test*, method = “spearman”). Data processing workflow can be found as a Jupyter-lab notebook at https://github.com/SebasSaenz/Papers_wf as well as other R scripts for data wrangling and visualisation. The MGnify cow rumen v1.0 MAG catalogue collection can be found at http://ftp.ebi.ac.uk/pub/databases/metagenomics/mgnify_genomes/cow-rumen/v1.0/ and the Hungate 1000 catalogue of reference genomes from the rumen microbiome is available at https://genome.jgi.doe.gov/portal/TheHunmicrobiome/TheHunmicrobiome.info.html.

## 3. Results

### 3.1. Dataset description

A total of 5990 rumen microbiome genomes were collected from the MGnify genome resource (42) and the Hungate1000 collection (**Table S1**). In summary, 5578 of the collected genomes were metagenome-assembled genomes (MAGs) and 412 were bacterial and archaeal isolates. After checking the quality of the assemblies using CheckM2, 2930 genomes were classified as high-quality drafts, 3035 as medium quality drafts and zero as low-quality drafts. The remaining genomes were discarded because they had more than 10% contamination or completeness could not be calculated. Based on genome quality, 3038 genomes with ≥ 90% completeness were retained for further analysis. Briefly, the selected genomes had a mean 107,308 bp ± 274,286 N50, 0.901 ± 0.0263 coding density, 337 bp ± 27.8 gene length, 2,535,744 bp ± 794,061 genome size and 48.5 ± 8.6% GC content (**Table S2**). In terms of taxonomic classification, 62 genomes were classified as *Archaea* by GTDB-TK (29), while 2976 were classified as *Bacteria*. In addition, 22 phyla were identified across the dataset, of which 10 were represented by more than 20 genomes. Genomes classified as *Bacillota* were the most abundant in the dataset representing 50.8% of the total genomes, followed by *Bacteroidota* (32.4%), *Pseudomonadota* (3.7%), *Actinomycetota* (3.4%), *Cyanobacteriota* (2.5%) and *Methanobacteriota* (1.7%). Moreover, genomes identified as the genus *Prevotella* (8.4%) were the most prevalent, followed by *Cryptobacteroides* (6.6%), uncultured CAG-791 (3.3%), *Ruminococcus* group E (3.0%) and *Sodaliphilus* (2.9%).

### 3.2. Prevalence of defence mechanism

Antiviral defence systems are prevalent in the rumen microbiome. In total, 14,241 defence systems and 31,948 antiviral related genes, representing 114 unique system types, were identified across all genomes. Out of the 3038 genomes, 89% of them carry at least one antiviral defence system. Approximately, 99% of isolates harboured at least one system, compared to 87% of the MAGs (**Figure 1A**). On average, isolates had a larger genome (**Figure S1**), number of defence system families (p-value<2.2e-16, **Figure 1B**), number of defence systems (p-value<2.2e-16, **Figure 1C**) and defence systems density (p-value=1.051e-15, **Figure 1H**) compared with the MAGs. Similarly, 96% of archaeal genomes harboured at least one defence system, compared to 88% of bacterial genomes **(Figure 1D**, **Table S3)**. On average, *Archaea* seem to carry 4.9 systems per genome while *Bacteria* 4.7 (p-value=0.1875, **Figure 1F**). However, this number highly varies, being zero the minimum (3.2% archaeal and 11.1% bacterial genomes) up to 34 in the Ca. *Limimorpha*. Approximately 92% of archaeal and bacterial genomes carry between one to ten systems but the remaining genomes harbour between 11 to 34 and 11 to 19 systems in *Bacteria* and *Archaea* respectively **(Figure 2A)**. The density of defence systems per genome significantly differed between *Archaea* and *Bacteria* (p-value = 0.003752, **Figure 1G**), 2.12×10^-3^ and 1.75x^-3^, respectively. Additionally, after grouping the systems into families we found that the average number of unique antiviral defence families in archaeal and bacterial genomes is similar, 2.8 and 2.6 respectively (p-value = 0.1456, **Figure 1E**). The minimum number of families found was zero and 13 the maximum in the uncultured *Lachnospiraceae* CAG-603.

**Figure 1.**
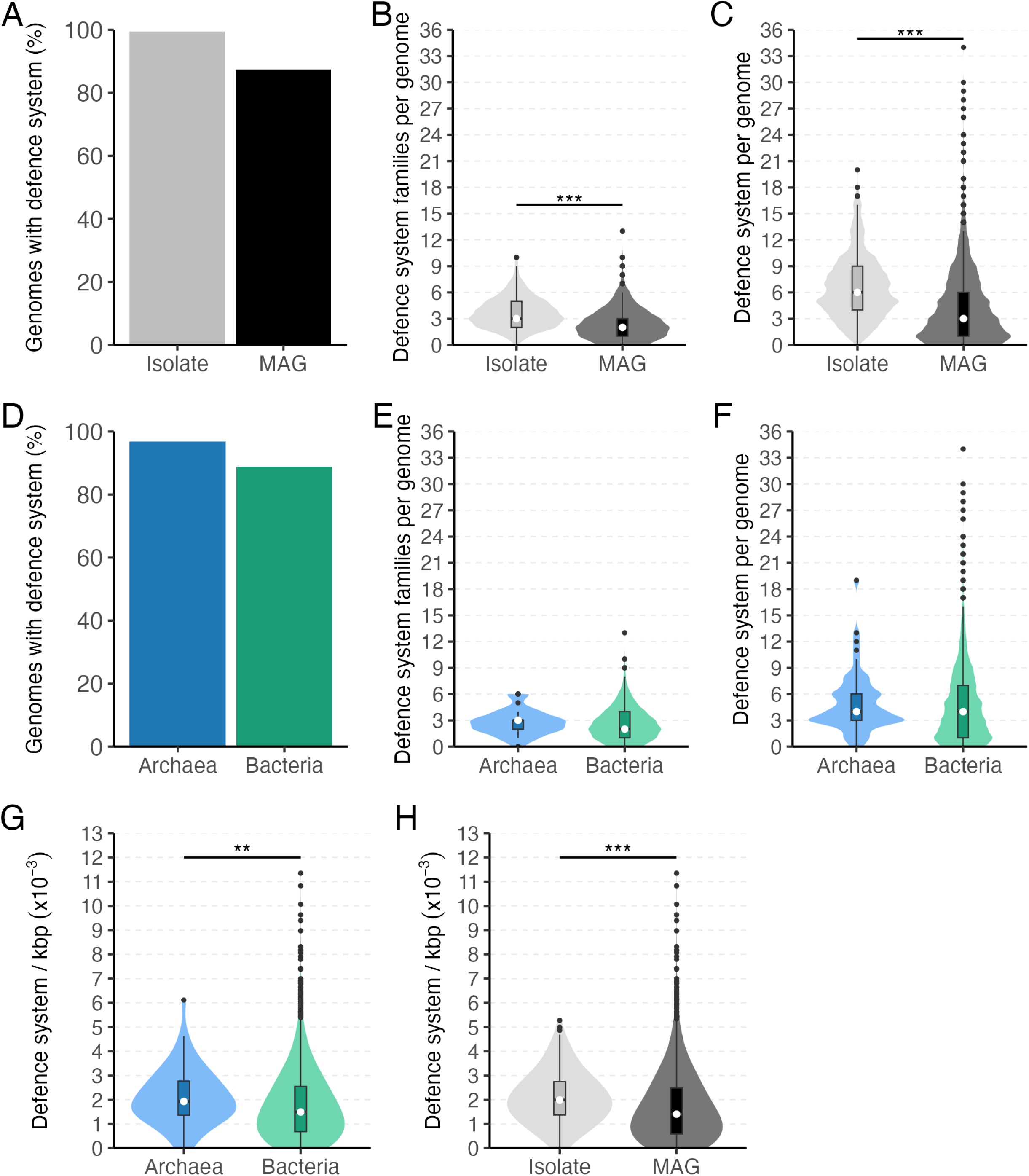
Prevalence of defence mechanisms in the rumen. A) the percentage of genomes with at least one defence mechanism, B) the number of defence systems per genome, C) the number of unique defence system families per genome in isolates and MAGs. D) The percentage of genomes with at least one defence mechanism, E) the number of defence systems per genome, F) the number of unique defence system families per genome in *Archaea* and *Bacteria*. The defence system density (per genome per kbp) in G) *Archaea* and *Bacteria* and H) isolate and MAGs

**Figure 2.**
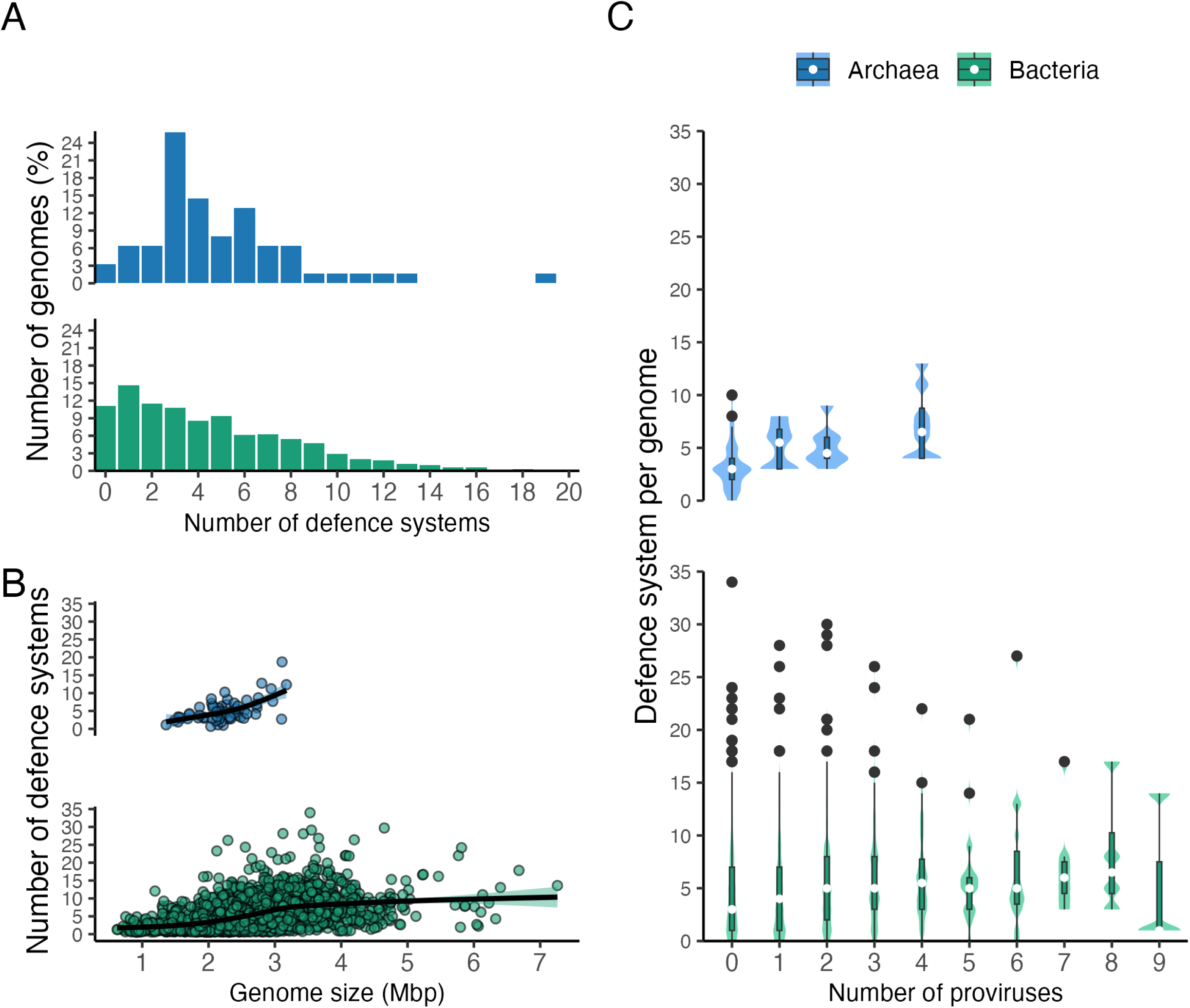
Distribution of the total number of antiviral systems per genome and their correlation with genomes size and prophage prevalence. A) the percentage of genomes carrying different number of defence systems, B) correlation between the number of defence system per genome and the genome size and C) correlation between the number of defence system per genome and the number of proviruses per genome.

When compared by phyla, between 78% and 100% of the genomes carry at least one defence system (**Figures S2**), being *Verrucomicrobiota* and *Fibrobacterota* the highest and *Pseudomonadota* the lowest. *Bacillota* and *Bacteroidota* phyla are among the most prevalent in the dataset, with 84% and 96% of the genomes exhibiting defensive mechanisms respectively. Similarly, in average *Verrucomicrobiota* and *Fibrobacterota* genomes have the higher number of defence systems and defence systems density (**Figure S2**).

We then used Virsorter2 to detect proviral sequences longer than 10,000 bp in the genomes and CheckV to evaluate their quality. The results were compared with the number of defence mechanisms found in each genome. Forty percent of the genomes harboured at least one provirus, being zero the lowest and 10 the highest in one bacterial genome. A positive correlation was observed between the average number of defence systems and the number of proviruses per genome in archaea (Spearman ⍴=0.575, p-value= 1.005e-06, **Figure 2C)**, however this was not observed in the bacterial genomes (Spearman ⍴=0.080, p-value= 1.04e-05). In contrast, the genome size positively correlates with the number of defence systems in archaeal (Spearman, ⍴=0.528, p-value=1.421e-05) and bacterial genomes (Spearman, ⍴=0.525, p-value < 2.2e-16, **Figure 2B**). The relation between the number of proviruses and the genome size has been previously evaluated as potential drivers of the number of antiviral systems in prokaryotic genomes and plasmids (23, 43).

### 3.3. Diversity and abundance of defence mechanism

To estimate the diversity of antiviral defence systems, all predicted systems were grouped into families using the metadata provided by PADLOC-DB. In total, 114 different system types were found across all genomes, and they were grouped in 49 families. All of them were found in bacterial genomes, whereas only 16 of them were found in the archaeal genomes. RM, Abi and cas system families were present in 72.1, 55.2 and 27.9 % of all genomes. Moreover, 28 of the families were present only in 1% of the genomes. In terms of relative abundance, systems belonging to RM were the most abundant in *Archaea* (51.1%) and *Bacteria* (44.5%) compared to the total systems found **(Figure 3A and 3B)**. However, not always the most abundant systems were equally present in both groups. For example, the abundance of the Abi system was 6.9 and 24.6% in *Archaea* and *Bacteria* respectively. Similarly, the family AVAST represented 5.6 and 0.4% of all systems found in *Archaea* and *Bacteria*. Additionally, the relative abundance of 34 of the total families was less than 1%.

**Figure 3.**
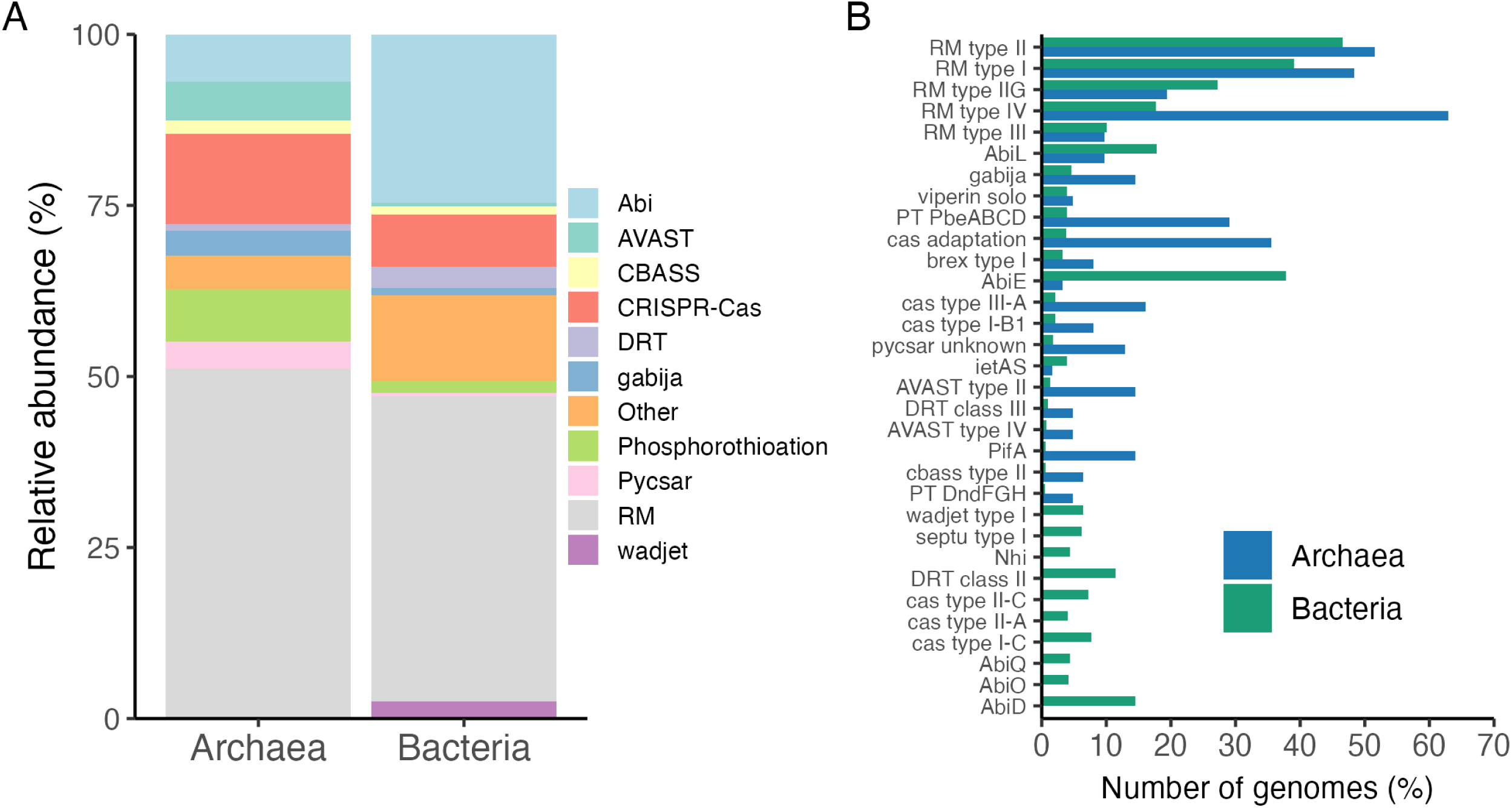
Abundance and diversity of defence mechanisms within *Bacteria* and *Archaea* in the rumen. A) The relative abundance of all defence systems grouped by families in *Bacteria* and *Archaea* and B) the percentage of genomes per defence system. Defence system families with a relative abundance < 1.5 % were grouped in the “other” category. The defence systems that were found in < 2% of genomes are not shown.

### 3.4. Defence mechanism in key members of the rumen

To corroborate if the antiviral defence systems are enriched in a specific taxon, we selected all genera represented by more than 10 genomes (n=57). The number of defence systems and families of the selected genera were compared. Both total numbers of systems and number of different families vary between genera. On average, these genomes (genera represented by at least 10 genomes) carried 4.6 defence systems and 2.6 defence system families. The uncultivated genus Ga6A1 (12.8) and Ca. *Limimorpha* (12.6) had the highest number of systems, followed by *Fibrobacter* (9.8) **(Figure 4B)**. Similar trends were observed when the number of families were compared **(Figure 4A)**. None of the genera were totally devoid of defence systems but *Enterousia* and several uncultivated bacteria had a low prevalence of systems. Additionally, we observed a positive correlation between the number of system and families, which indicates that *Bacteria* or *Archaea* carrying several defence systems will also harbour more defence system families (**Figure S3**)

**Figure 4.**
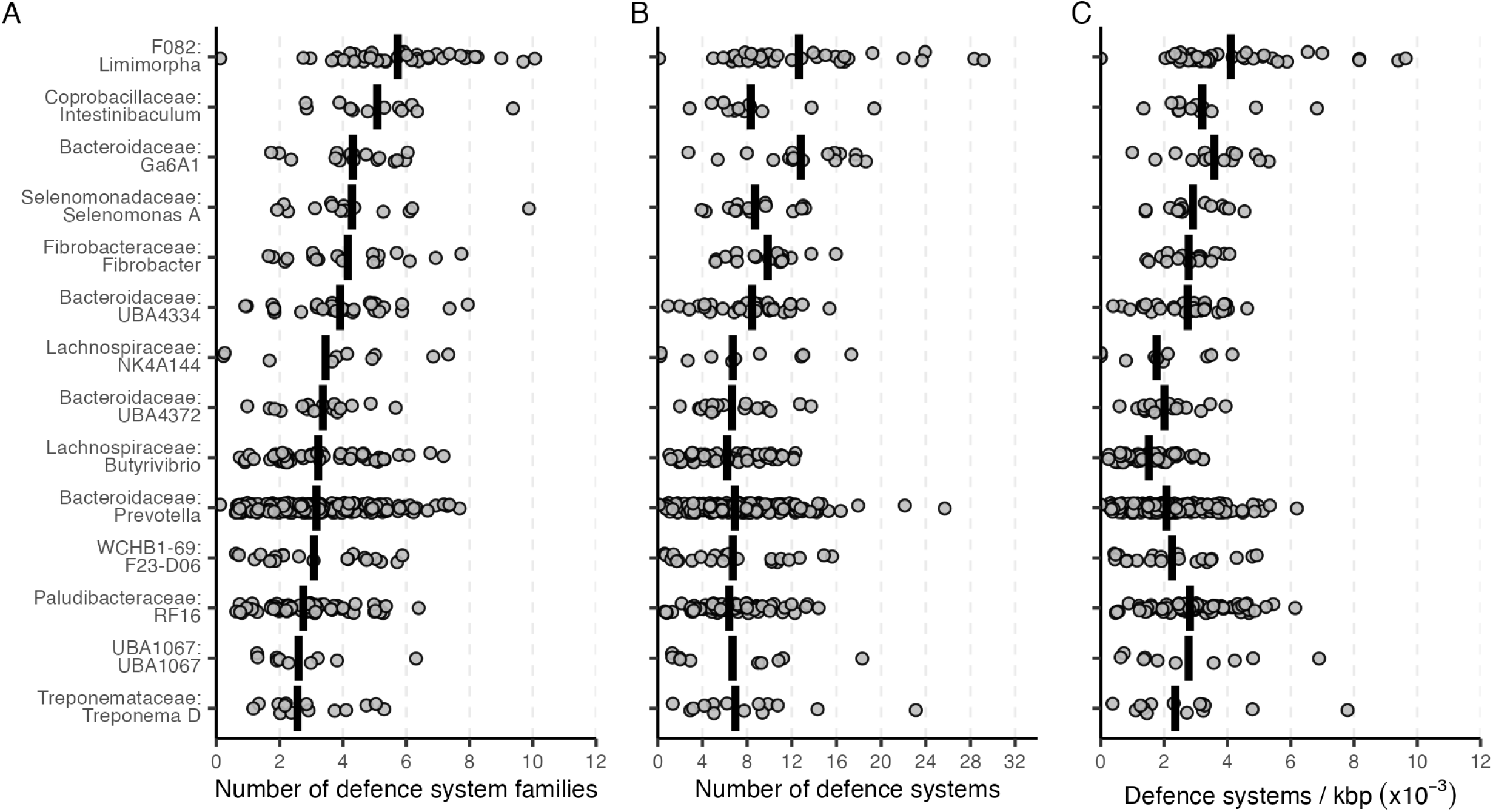
Number of defence systems in different genera from the rumen. A) Number of defence system families, B) number of defence systems and C) defence system density (per genome per kbp) found in the top 14 genera with the highest average number of defence systems per genome. The vertical bar represents the average number of defence systems and system families. Genera with at least 10 genomes are shown. The labels in the x-axis represent the Family:Genus.

Additionally, the diversity of defence system and system families varies across the dominant members of the rumen (**Figure 5A**, **S4**). For example, between 37-97% of genomes of the genera *Prevotella*, Ca. *Limimorpha*, *Butyrivibrio* and *Ruminococcus* carry RM and Abi systems (**Figure 5B**). However, 22, 31, 24, and 23 different defence system families are present in less than 20% of the same genomes respectively. The Ca. *Limimorpha* seems to be an example of a genus carrying a core group of defence genes, because at least 50% of its genomes harboured RM, Abi, Pycsar, cas and DRT systems (**Figure 5B**).

**Figure 5.**
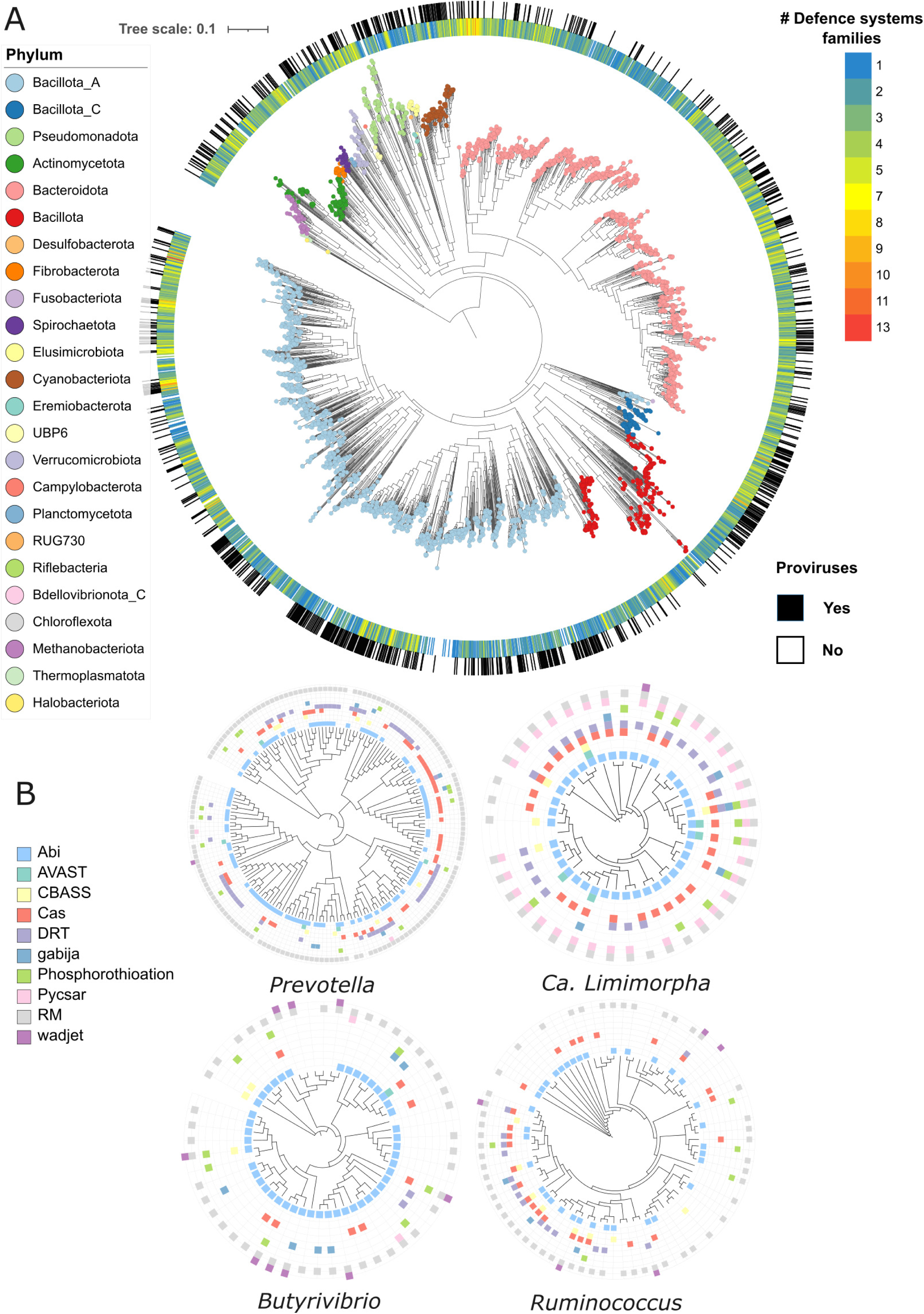
Distribution of defence mechanisms across the rumen. A) Phylogenetic tree of the rumen microbiome (3038 genomes) based on 400 universal markers. b). Phylogenetic tree based on core genes of four members of the rumen microbiome: *Prevotella*, Ca. *Limimorpha*, *Butyrivibrio* and *Ruminococcus*.

### 3.5. CRISPR-Cas arrays and phage interaction

Cas proteins and CRISPR arrays have been reported across complete or draft prokaryotic genomes from different environments (43, 44). However, the accurate automated prediction of CRISPR-Cas loci is challenging due to the variability in genome quality and the ambiguous prediction of the system subtypes. Therefore, we used CRISPRCASTyper to identify complete loci and the system subtype based on both Cas genes and CRISPR repeats in the rumen genomes. In summary, 791 CRISPR-Cas complete locus were found in 668 genomes. This indicates that most of these genomes carry only one complete system. In total, 21.9% of all 3038 rumen microbiome genomes harbour at least one CRISPR-Cas complete locus. When compared, more bacterial genomes seem to carry more complete systems than archaeal genomes, 22.1 and 14.5 % respectively **(Figure 6A)**. The phylum with the highest prevalence of CRISPR-Cas was the uncultured UBP6 (42% of genomes), followed by *Actinomycetota* (31%), *Bacteroidota* (26%) and *Verrucomicrobiota* (26%) **(Figure 6B)**. Interestingly, of the phyla represented by at least 50 genomes, only *Cyanobacteriota* was almost devoid of complete CRISPR-Cas. In this case, one genome out of 77 carried an array. In total, 21 CRISPR-Cas subtypes were predicted across all genomes and among them the subtype I-C was the most abundant (21.9%), followed by II-C (15.1%), II-A (13.5%) and I-E (10.3%). The number of subtypes found per phyla greatly varied, being 15 the maximum in *Bacillota* followed by 13 in *Bacteroidota*. However, in the phylum *Spirochaetota*, *Eremiobacterota*, *Elusimicrobiota*, and *Desulfobacterota* only one subtype was found in all genomes **(Figure 6C)**.

**Figure 6.**
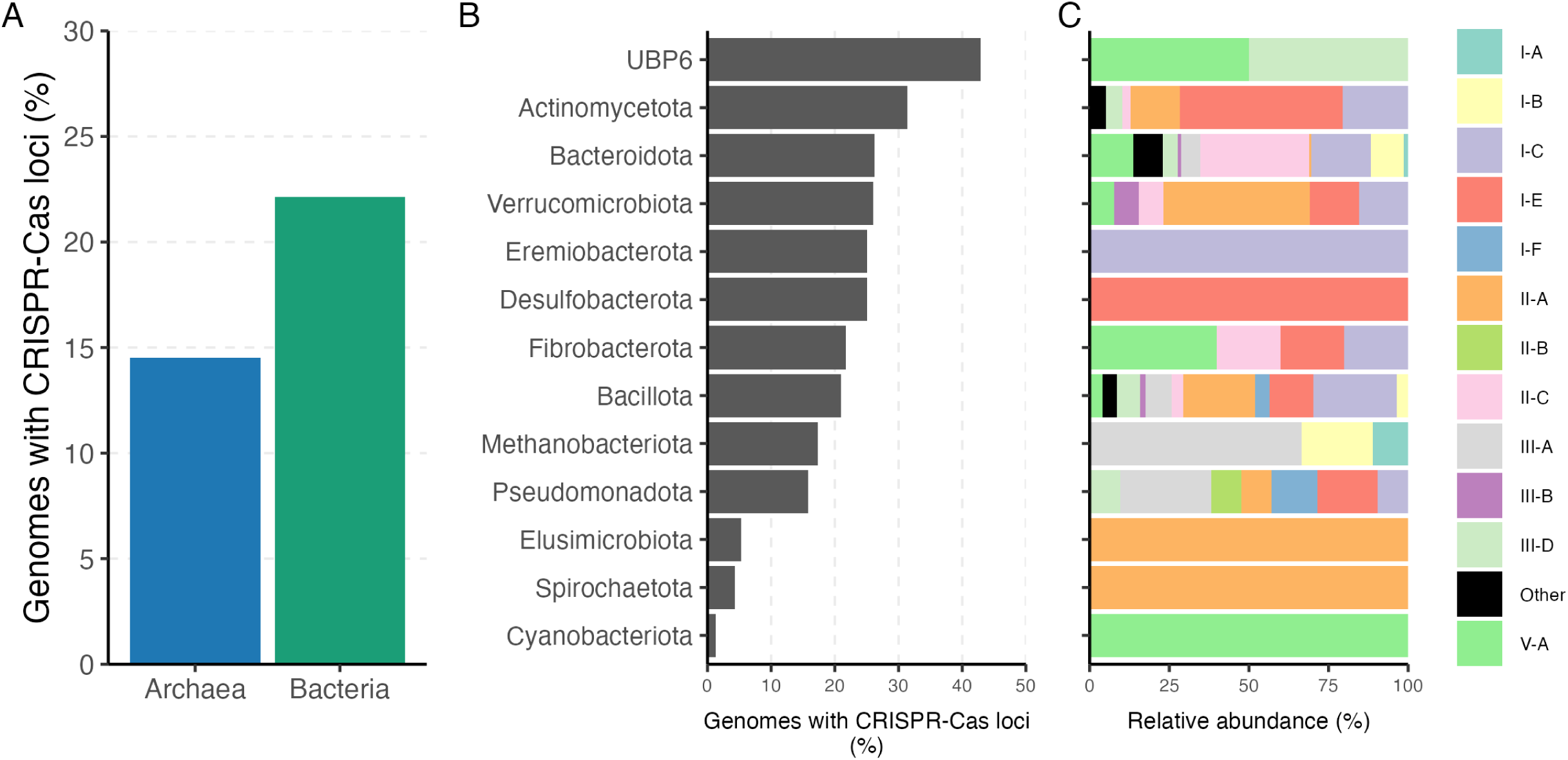
Abundance and diversity of CRISPR-Cas systems within bacteria and archaea in the rumen. A) Percentage of genomes harbouring at least one complete CRISPR-Cas loci, B) percentage of genomes by phylum harbouring at least one complete CRISPR-Cas loci, and C) the relative abundance of CRISPR-Cas system types in each phylum.

All CRISPR spacers predicted by CRISPRCASTyper were collected, including those from incomplete CRISPR-Cas locus. In total, 27,717 spacers were found in 845 genomes, ranging from one to one hundred spacers per array. All collected spacers were aligned against a rumen virome database (RVD) using blast, containing 397,180 species-level viral operational taxonomic units (vOTUs). The vOTUs were mined from 975 public rumen metagenomes, including samples from four different continents and 13 animal species. In total, 30.2% of the spacers matched against 4986 vOTUs, which could indicate a previous history of phage-bacterial and phage-archaeal infection (**Figure 7**, **Table S4**). Most of the interactions were found between members of the phylum *Bacillota*, *Bacteroidota* and *Actinomycetota* and different vOTUs. For example, two uncultured species from the *Firmicutes* group A have a history of interaction with 53 and 52 different vOTUs, which was the maximum found (**Table S5**). Similarly, the spacers of the archaea *Methanobrevibacter* and bacteria *Prevotella* matched with 49 and 39 different vOTUs, respectively. Additionally, approximately 7% of the collected phages showed intra-species host specificity, while 2.6% intra-genus host specificity (**Table S6**). Only 1.8% of the vOTUs that matched with the spacers have been classified as members of the *Siphoviridae* and *Myoviridae* families, while the rest remain unclassified.

**Figure 7.**
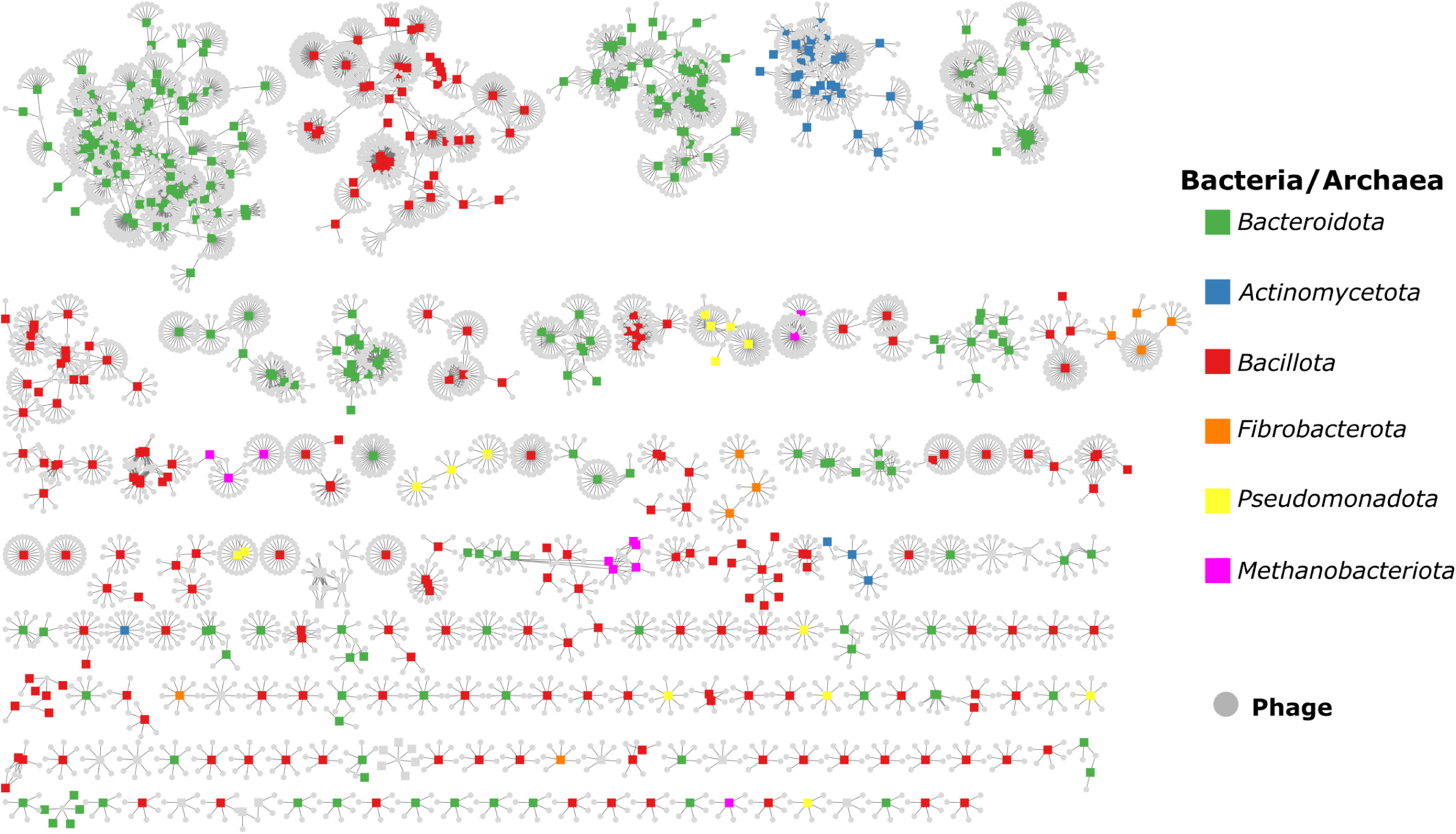
Interaction between rumen phages and their host. The host-virus interaction network shows the match between the identified CRISPR spacers from this study and species-level viral operational taxonomic units (vOTU) obtained from the global rumen virome database (RVD). Grey circles indicate a vOTU while squares represent bacterial/archaeal genomes coloured based on their phylum.

## 4. Discussion

It is estimated that phages are 10-times more abundant than bacteria and their role in shaping microbial communities, influencing genetic diversity, and participating in biogeochemical cycles is a crucial aspect of the Earth’s ecosystems (45, 46). By exerting evolutionary pressure on host populations, phages drive the evolution of their hosts. In the rumen, phages are diverse and infect the core microbiome, including *Prevotella*, *Ruminococcus*, *Fibrobacter*, *Butyrivibrio*, and Methanobrevibacter (13, 47). This is especially important because the arm-race between phages and host can modulate the functional traits in the rumen (14). This arm-race is driven by defence systems in prokaryotes and by anti-defence proteins in phages and other MGE (18, 19). In the rumen we found that defence systems are diverse, mainly associated to innate immunity and dominated by RM, abortive systems and cas proteins. The high prevalence of defence systems with a broad genetic organisation suggests that selective pressure has been exerted by phages and other MGEs in the rumen. However, this pressure also triggers the evolution of antidefence encoded in phages. For example, proteins that inhibit CRISPR-Cas, Gabija, Thoeris and Hachiman systems are present in phages that infect taxonomically diverse bacterial species (19, 48, 49). The rumen is not the exception as it has been found that rumen phages carry anti-RM system, which represents a counter-defence mechanism of the most abundant prokaryotic defence (13). The emergence of different strategies to evade defence systems in phages and other MGEs could drive the rapid acquisition of diverse and novel defence systems in prokaryotes. This arm-race extends even between viruses, as phages are reservoir of antiviral systems which can inhibit competing viruses (50, 51).

The high variability of defence systems and defence system families across the genomes from different environments, including the rumen, could back the ‘pan-immune system’ model (20). This model suggests that genetically closely related strains could share defence systems via horizontal gene transfer, which indicates that the immunity could be driven by the collection of systems in the community. Mobile genetic elements (MGEs) play a main role in the mobilization of defence systems because they often carry defence genes, even more frequently compared to chromosomes (52, 53). Moreover, defence system seen to organize in defence island, accompanied by MGE, from which novel defence genetic architecture could emerge (21, 52). In the rumen, the pan-immunity could be driven mainly by the innate response, as CRISPR-Cas is only present in approximately 21% of the rumen genomes, but they have been shown to be important in the evolutionary dynamic between phage and their host in the rumen (14). This seems to be common also in other environments like cheese and silage-associated bacterial communities (54, 55). The genomes analysed in the present study harbouring CRISPR-Cas systems showed a history of infection with phages mined from rumen samples. Some of this phages could infect not only several genomes but also species and genera, which has been previously reported in samples from ruminants (56), the human gut (57) and the ocean. A broad host range could possibly have arisen through adaptation to different receptors by mutations, which is also considered as an advantage for the arm-race without acquiring anti-defence systems (58). Such polyvalent phages can influence the rumen metabolic traits at the cellular and microbiome level. For example, our data showed that *Prevotella* and *Methanobrevibacter* with a wide repertoire of defence mechanism, involved in fiber degradation and methanogenesis, have a broad interaction with phages (59, 60). This was consistent with previous reports on the modulating role of the rumen viral community in key metabolic traits like nutrient recycling, fiber degradation and methanogenesis. (14, 61). This shows that even though in low proportion the adaptive immunity can participate in the response of specific infections that can modulate the microbial and animal metabolism.

Moreover, the present study adds to the growing evidence that highlights the prevalence and significance of antiviral defence systems in prokaryotic communities. We show that defence mechanisms in the rumen microbiome are prevalent, with almost 90% of the bacterial and archaeal genomes carrying at least one system. A similar number of defence systems has been observed within genomes from the NCBI RefSeq (23) and MAGs from soil and human gut (**Figure S5**) (52). Additionally, the number of systems per genome from different environments seem to follow a binomial distribution, where most genomes carry between 2-5 systems. We observed a similar distribution within the families of the defence systems in the rumen. The combination of different systems can provided a broader resistance and resistance range (62, 63) and the widespread distribution of these systems suggests a crucial role in maintaining microbial diversity and ecosystem stability.

Also, our findings further reinforce the notion that defence mechanisms are diverse but just a few are dominant in the rumen environment. Consistently, RM, Abi and Cas are the most abundant defence system families across different genomes and environments (52, 64). At least in the rumen these three families represent approximately 70% of all defence systems. Furthermore, 28 out of the total 49 families found in the rumen were present only in 1% of the genomes. This shows that these systems are rare, but they are still present in diverse genomes. Such numbers have been observed in a collection of 21000 RefSeq genomes, which also suggests that the diversity of antiviral systems is bigger than previously thought (23).

Different environmental and genetic factors can contribute to the selection of genes (65, 66). Previous studies have shown that bacterial and phage genome, and plasmid size seems to drive the acquisition of defence mechanisms (23, 43, 67). We also found that this is true for bacteria and archaea in the rumen, as the number of defence systems positively correlated with genome size. Genomes would tend to be larger when defence systems such as CRISPR-Cas are present due to their size, and the same could apply when multiple small defence systems are recruited in the same genome or defence islands (52). On the other hand, the number of prophages in the genome had a low or positive correlation with the number of defence systems. We were expecting a negative correlation between the number of defence systems and prophages, as more systems can provide a broader resistance, but it was not the case in our samples. However, other studies have found that the abundance of CRISPR-Cas systems can be linked with the viral abundance (68).

It is relevant to point that our study has a few limitations. Firstly, most of the used genomes in the study are product of MAG binning, which is known to poorly reconstruct closely related strains (69). This is especially important, because it could limit the detection of defence systems due to the variations of defensome composition between strains (21). Additionally, short-sequencing may not provide accurate genome-specific signals, low abundance, and could affect the sensitivity/specificity trade-off of the single-gene or multi-gene system detection specially in large CRISPR arrays (22)

. Secondly, the unbalanced number of isolates and MAGs shows that the rumen microbiome is still largely unculturable. In this way the description of the defensome is mostly computational dependant. Also, isolates tend to harbour more defence system, defence system families and a higher defence system density, which can be associated with the genome completes and low fragmentation. The accumulation of more defence mechanism in isolates compared to MAGs can be mainly associates to the methods used to recover them and not their lifestyle or role in the microbial community. However, a larger portion of isolates need to be collected to draw robust conclusion. Besides these limitations, we believe that this targeted data analysis is a starting point to expand the search for defence system in the rumen using state of the art techniques such as long sequencing and high throughput culturomics.

## 5. Conclusion

Antiviral defence systems are prevalent and diverse in the rumen, but only a few are dominant, indicating that many systems are rarely present. However, the collection of systems throughout the rumen may represent a pool of mechanisms that can be shared by different members of the community. This could support the ’pan-immune system’ model, which appears to be common across different environments.

## Ethics approval and consent to participate

Not applicable

## Availability of data and materials

Data processing workflow can be found as a Jupyter-lab notebook at https://github.com/SebasSaenz/Papers_wf as well as other R scripts for data wrangling and visualisation. The MGnify cow rumen v1.0 MAG catalogue collection can be found at http://ftp.ebi.ac.uk/pub/databases/metagenomics/mgnify_genomes/cow-rumen/v1.0/ and the Hungate 1000 catalogue of reference genomes from the rumen microbiome is available at https://genome.jgi.doe.gov/portal/TheHunmicrobiome/TheHunmicrobiome.info.html.

## Authors’ contributions

JSS collected and analysed the data, prepared the figures and wrote the manuscript. BRG contributed to the analysis of the data. JS contributed to the discussion of the data, supervision and funding. All authors revised and approved the final version of the manuscript.

## Acknowledgements

The authors acknowledge support by the High Performance and Cloud Computing Group at the Zentrum für Datenverarbeitung of the University of Tübingen, the state of Baden-Württemberg through bwHPC and the German Research Foundation (DFG) through grant no INST 37/935-1 FUGG.

